# Spt5 interacts genetically with Myc and is limiting for brain tumor growth in Drosophila

**DOI:** 10.1101/2023.04.14.536839

**Authors:** Julia Hofstetter, Ayoola Ogunleye, André Kutschke, Lisa Marie Buchholz, Elmar Wolf, Thomas Raabe, Peter Gallant

## Abstract

The transcription factor SPT5 physically interacts with MYC oncoproteins and is essential for efficient transcriptional activation of MYC targets in cultured cells. Here we use *Drosophila* to address the relevance of this interaction in a living organism. Spt5 displays moderate synergy with Myc in fast proliferating young imaginal disc cells. During later development, Spt5-knockdown has no detectable consequences on its own, but strongly enhances eye defects caused by Myc-overexpression. Similarly, Spt5-knockdown in larval type 2 neuroblasts has only mild effects on brain development and survival of control flies, but dramatically shrinks the volumes of experimentally induced neuroblast tumors and significantly extends the lifespan of tumor-bearing animals. This beneficial effect is still observed when Spt5 is knocked down systemically and after tumor initiation, highlighting SPT5 as a potential drug target in human oncology.

## Introduction

Expression of MYC oncogenes is deregulated in most human tumors. Up to 28 % of all tumors exhibit gene amplification of one of the MYC isoforms (MYCN, MYCL or MYC), defining MYC genes as the most frequently amplified oncogene family across human cancers (Schaub *et al*, 2018). Indeed, MYC is a crucial driver of tumorigenesis as demonstrated by mouse experiments involving MYC-overexpression (Adams *et al*, 1985; Kortlever *et al*, 2017), genetic depletion of endogenous (Sansom *et al*, 2007; Walz *et al*, 2014) or exogenous MYC (Jain *et al*, 2002), and expression of a dominant-negative variant of MYC (Soucek *et al*, 2008). MYC can therefore be considered a priority target for cancer therapy (Dang, 2012). At the same time, it is very challenging to target MYC directly, because it lacks enzymatic activity and probably pockets for small molecules (Nair & Burley, 2003). Instead, it seems possible to identify binding partners which the oncogenic function of MYC is fully dependent on, and to target them, for example the histone-methyl-transferase adapter protein WDR5 (Lorenzin *et al*, 2016; Thomas *et al*, 2015). In recent years, several additional MYC binding partners were identified by proteomic approaches, and MYC was shown to partake in multiple nuclear protein complexes (Baluapuri *et al*, 2019; Buchel *et al*, 2017; Dingar *et al*, 2018; Kalkat *et al*, 2018; Koch *et al*, 2007). To be considered as suitable for pharmaceutical targeting, such MYC binding partners should be (i) essential for MYC-driven oncogenic growth and (ii) dispensable for the integrity and proliferation of healthy tissue. The former is relatively easy to analyze systematically in transplantation-based murine tumor models (Vo *et al*, 2016), but the latter is very elaborate and expensive to study in mice. We therefore started to develop a *Drosophila* model to (i) validate the genetic interaction between MYC and its binding partners in vivo, and (ii) to estimate effects on healthy tissue of animals and thus the potential therapeutic window.

The *Drosophila* genome encodes a single MYC homolog that accomplishes the functions of its vertebrate counterparts in normal cells, and it also acts as an oncogene in *Drosophila* tumor models. Here, we focused on a brain tumor model derived from neural stem cells (type II neuroblasts = NB II), which allows to study proliferation and tumorigenesis during brain development. Briefly, NB II produce intermediate neural progenitors (INPs) with a restricted proliferation potential, which in turn generate ganglion mother cells as the precursors of neurons and glia cells (Homem & Knoblich, 2012). NB II express the cell fate determinant Brain tumor (Brat) and pass it on to their progeny (Bello *et al*, 2006; Betschinger *et al*, 2006; Lee *et al*, 2006). In case of brat mutations, INPs acquire NB II characteristics, resulting in large transplantable tumors (Bowman *et al*, 2008; Hakes & Brand, 2019; Janssens *et al*, 2014; Komori *et al*, 2018; Xiao *et al*, 2012). Brat belongs to the TRIM-NHL family of proteins, which regulate gene expression by reducing translation and causing degradation of multiple mRNAs (Connacher & Goldstrohm, 2021; Tocchini & Ciosk, 2015). Brat targets many mRNAs involved in NB self-renewal, including Myc (Betschinger *et al*., 2006; Loedige *et al*, 2015). We exploited this tumor model to address the potential for interfering with tumor formation by targeting Myc interaction partners.

As a proof of the target validation concept, we chose the MYC binding partner SPT5. First, SPT5 was detected as a binding partner of both MYC (Baluapuri *et al*., 2019) and MYCN (Buchel *et al*., 2017), indicating that the interaction between MYC proteins and SPT5 is evolutionary conserved. Second, recombinantly expressed MYC and SPT5 build stable dimeric complexes in vitro, demonstrating their direct interaction (Baluapuri *et al*., 2019). Third, SPT5 is essential for MYC-mediated transcriptional activation, which is considered a key oncogenic function of MYC (Baluapuri *et al*., 2019). A function of SPT5 in transcription was already evident upon its initial discovery in a pioneer genetic screen by Winston and colleagues in yeast. Several Suppressors of Ty (SPT) genes, including SPT5, were discovered, since their mutation reactivated the transcription of an auxotrophy gene that was silenced by proximal insertion of a Ty transposon (Winston *et al*, 1984). Subsequent work demonstrated direct interaction of SPT5 with SPT4 in yeast (Hartzog *et al*, 1998; Swanson *et al*, 1991) and the function of the mammalian SPT4/5 complex as a pausing factor named DSIF (DRB sensitivity inducing factor) (Wada *et al*, 1998). SPT5 binds RNA Polymerase II (RNAPII) and promotes transcriptional elongation and termination (Cortazar *et al*, 2019; Fong *et al*, 2022; Henriques *et al*, 2018; Hu *et al*, 2021; Parua *et al*, 2018; Parua *et al*, 2020; Shetty *et al*, 2017) and RNAPII processivity (Fitz *et al*, 2018) by binding to its DNA exit region, facilitating re-winding of upstream DNA and preventing aberrant back-tracking of RNAPII (Bernecky *et al*, 2017; Ehara *et al*, 2017). SPT5 homologues are found in all domains of life. SPT5 shares the N-terminal (NGN) and one KOW domain with its bacterial homolog NusG, but the eukaryotic protein contains several copies of the KOW domain and additional N- and C-terminal sequences (Yakhnin & Babitzke, 2014). While SPT5 is an essential protein, its interaction with MYC could indicate that tumor cells are more dependent on the full function of SPT5 than un-transformed cells. Here, we explored the functional interaction between Myc and Spt5 in vivo in *Drosophila* and analyzed the consequences of Spt5 depletion in brain tumors induced by brat-knockdown. We demonstrate a clear genetic interaction between Myc and Spt5 in developing eyes and a functional role of Spt5 in neuroblast proliferation. Strikingly, systemic knockdown of Spt5 from late larval stages onwards inhibits tumorigenesis, but is tolerated by normal tissue and massively extends the life span of tumor prone flies. This demonstrates not only that SPT5 is an attractive candidate for targeting MYC-mediated oncogenic growth, but also suggests that inhibition of an essential process, such as transcription, could open a therapeutic window in tumor treatment.

## Results

### Genetic interaction of Spt5 and Myc in Drosophila

The *Drosophila* genome encodes a single SPT5 homolog (Kaplan *et al*, 2000), which is 50% homologous to human SPT5 and contains all identified protein domains (Fig. 1A). To investigate its genetic interaction with Myc we focused on adult eye phenotypes, which are known to be highly sensitive to alterations in Myc levels. Myc-overexpression in post-mitotic cells of this tissue (using GMR-GAL4; Fig. 1B) induced excessive growth and apoptosis, resulting in oversized and aberrantly shaped adult eyes (Montero et al. 2008, Steiger et al. 2008; Figs. 1C, S1A-B). siRNA-mediated Spt5-knockdown had no discernible effect on control eyes (Figs. 1C-D, S1C). Knockdown of Spt5 in the Myc-overexpression context however dramatically altered eye morphology leading to a glassy surface, suggestive of apoptotic cell loss and ensuing fusion of neighboring ommatidia (Figs. 1C, S1D). This phenotype was fully penetrant and accompanied by a reduction in overall eye size (Fig. 1D). Importantly, this effect was not due to experimental off-target artefacts, since it was completely rescued by expression of a mutated Spt5 cDNA that codes for wild type Spt5 protein but is not recognized by the siRNA (Figs. 1C, S1G-H). Overexpression of the siRNA-resistant Spt5 itself showed effects neither in control nor in Myc-overexpressing flies (Figs. 1C, S1E-F). Together, these observations demonstrate that Myc and Spt5 functionally interact and suggest that the transcriptional program activated by excessive Myc levels is critically dependent on Spt5.

**Figure 1.**
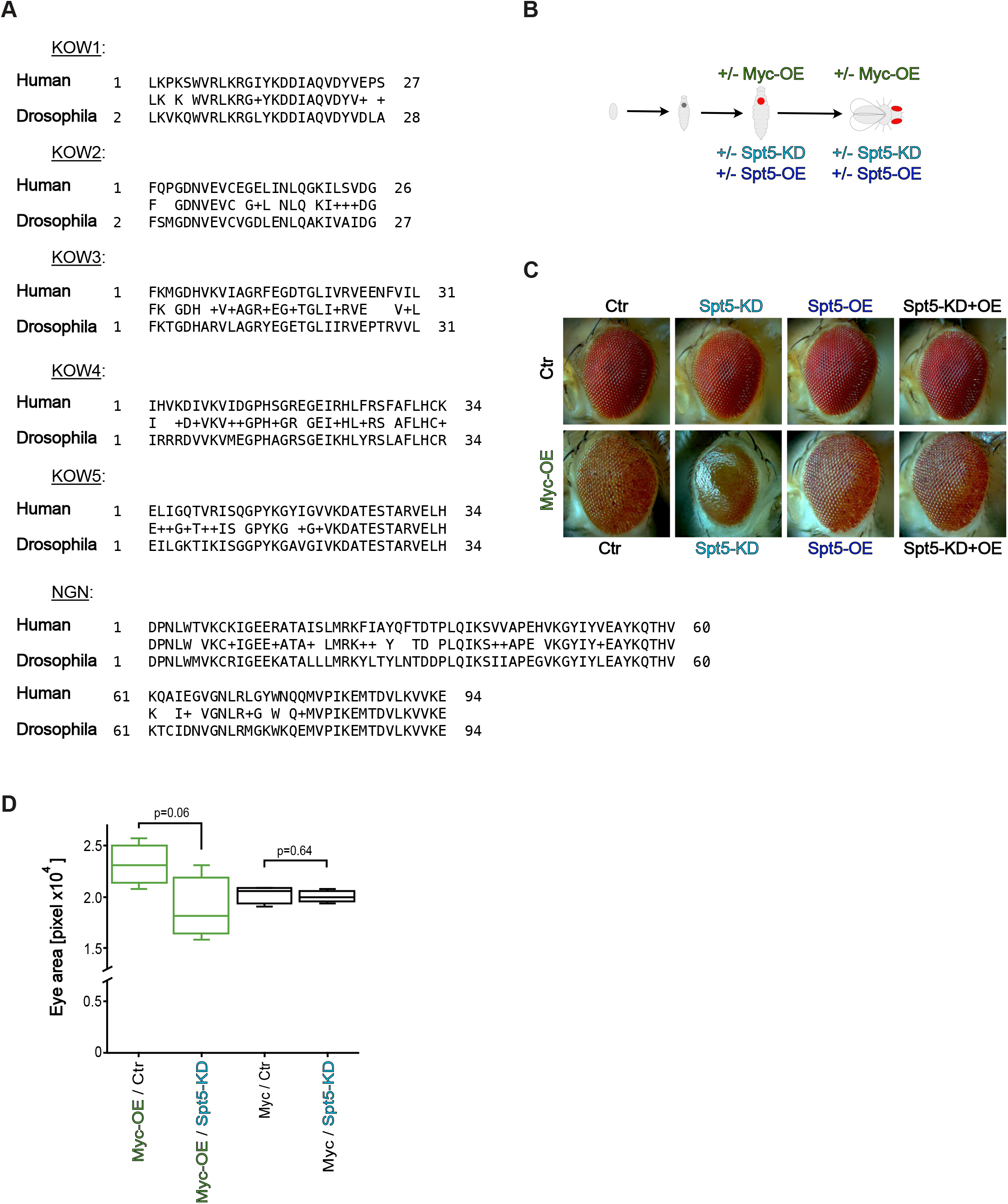
Genetic interaction of Spt5 with overexpressed Myc. ***A***, alignment of *Drosophila melanogaster* and human Spt5 proteins over all identified domains. ***B***, scheme depicting GMR-GAL4 dependent transgene expression in differentiating eye imaginal disc cells from the second half of the 3^rd^ larval instar onward. GAL4 activates expression of a Myc cDNA and/or an Spt5 siRNA and/or an Spt5 cDNA (coding for wildtype Spt5 protein, but refractory to siSpt5). ***C***, representative pictures of adult eyes of the indicated genotypes. ***D***, quantification of the eye areas from control (black) or Myc-overexpressing (green) flies. Median adult eye size from 4 independent flies each. P value was calculated using unpaired Student’s t-test.

Next, we addressed the organismal role of Spt5 during development. As described for yeast, Spt5 is an essential gene and Spt5 homozygous mutant flies do not reach adulthood (Mahoney *et al*, 2006). Spt5 heterozygotes were largely normal, except for a small but statistically significant reduction in adult body weight (Fig. 2A). Such a weight defect was also described for hypomorphic *Myc^P0^* mutants, which additionally show a slight delay in development (Johnston et al. 1999). The combination of both mutations did not affect the Myc-dependent developmental delay (Fig. S2A), but resulted in a synergistic weight loss (BLISS score 14, SynergyFinder; Fig. 2A). In addition, such doubly mutant flies showed deformed eyes (not shown). Such an eye defect was not observed in either single mutant alone, but had previously been described as a typical manifestation of the strong genetic interaction between Myc and its partner RUVBL1/pontin (Bellosta *et al*, 2005).

**Figure 2.**
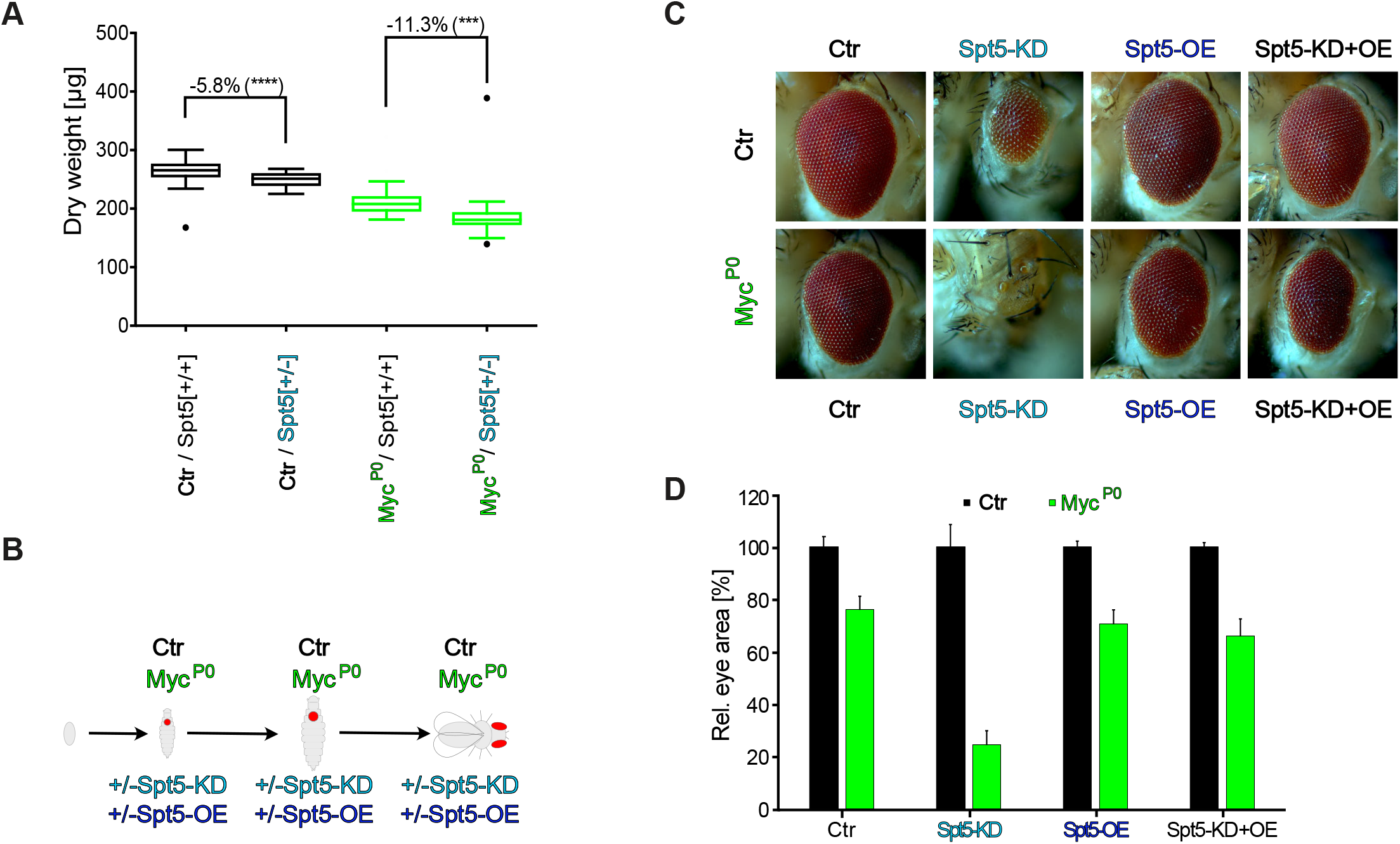
Genetic interaction of Spt5 with a hypomorphic Myc-mutant. ***A***, median dry weight of adult males (n=6-109) with or without Spt5-knockdown, in a Myc^wildtype^ (“Ctr”, black) or Myc^P0^ (green) background. P values were calculated using an unpaired Student’s t-test. ***B***, scheme illustrating the genetic manipulation: a ubiquitously expressed Myc cDNA was eliminated specifically in eye imaginal discs throughout larval development, thereby exposing the hypomorphic Myc^P0^ allele or Myc^wildtype^ (“Ctr”) while simultaneously driving Spt5-overexpression and/or -knockdown (see Methods). ***C***, representative pictures of adult eyes. ***D***, quantification of eye areas, normalized in each case to the area of the matching Myc^wildtype^ flies (“Ctr”, black); n=8 flies per genotype.

The synergy between Spt5 and Myc in proliferating cells became even more evident when Spt5 and Myc levels were reduced specifically in developing eye imaginal discs (Fig. 2B). In line with earlier publications, partial loss of Myc in this system impaired growth of eye imaginal disc cells and resulted in smaller adult eyes made up of smaller ommatidia (Figs.2C-D, S2B-I; Bellosta *et al*., 2005). Combination of the partial loss of Myc with Spt5-knockdown showed clear synergy (BLISS score 16, SynergyFinder) and nearly eliminated eye development. These observations confirm a functional collaboration between Spt5 and Myc in the control of cellular growth and proliferation.

### Effect of Spt5 on NB II-tumor development

Having confirmed the importance of Spt5 for Myc-dependent physiological processes, we set out to explore the role of Spt5 in brain tumors that were induced by knockdown of the tumor suppressor brat specifically in larval NB II. The adult brains of brat-knockdown animals are enlarged with a massive increase of cell number in the cortex region and a complete disruption of neuropil structures (Figs. 3A-B). In contrast, knockdown of Spt5 in NB II had only minor effects on adult brain structures, e.g. resulting in a ventral opening of the ellipsoid body of the central complex, which is one descendant of NB II cell lineages. Simultaneous knockdown of Spt5 and brat abrogated the overgrowth phenotype and largely restored normal brain structures (Fig. 3B). To quantify this effect, we expressed luciferase in the cells experiencing brat-knockdown. Luminometry of total lysates from young adults confirmed the strong growth-suppressive effect of Spt5-knockdown specifically in tumorous animals as opposed to control animals; expression of the siSpt5-insensitive Spt5 transgene abrogated the effects of siSpt5, demonstrating its specificity (Fig. 3C). Consistent with these findings, brat-knockdown led to a massive expansion of NB II cell lineages, which was largely abolished by simultaneous Spt5-knockdown (Fig. S3A).

**Figure 3.**
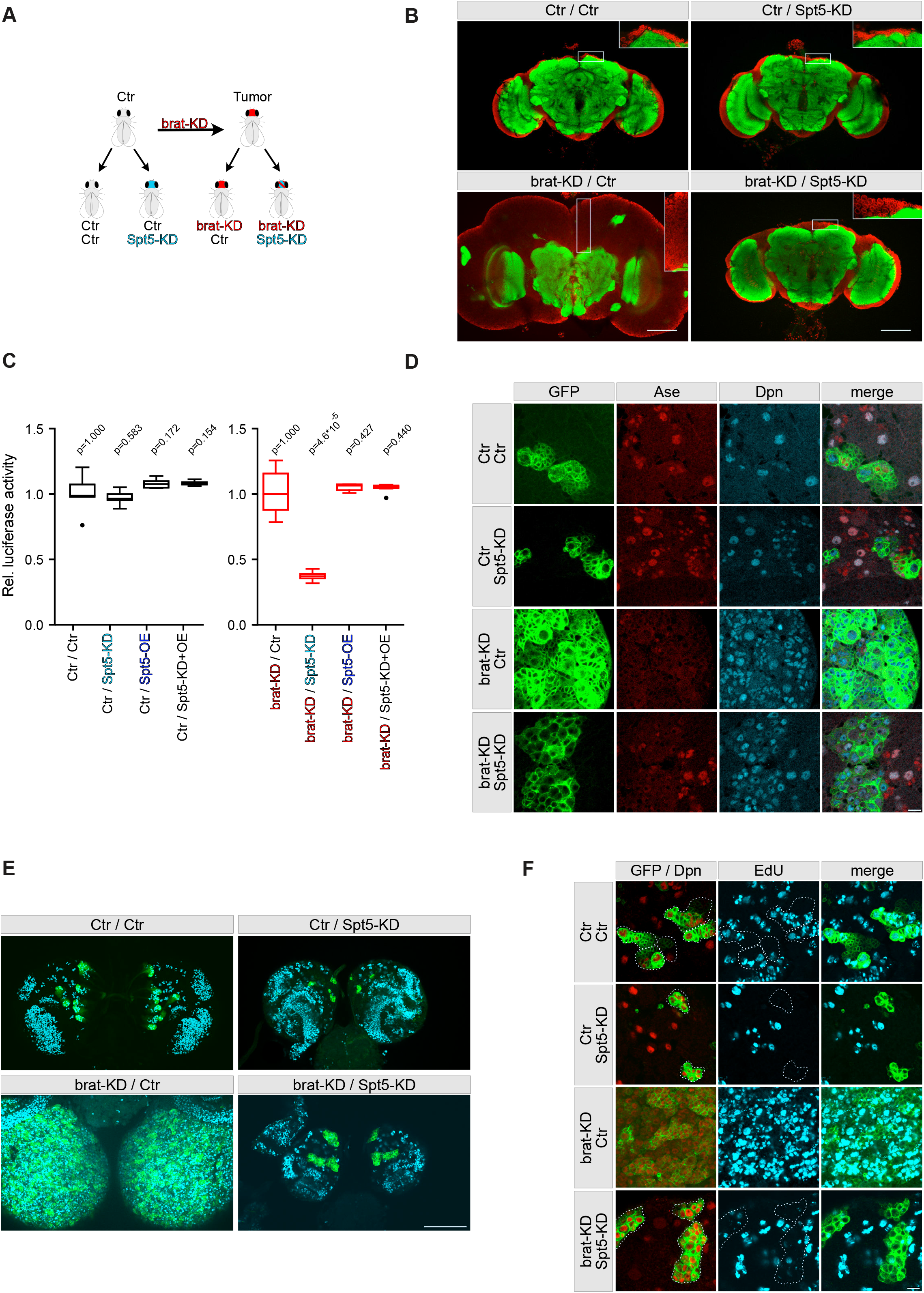
Spt5-knockdown reduces growth of brat-depleted tumors. ***A***, scheme of the NB II tumor model, showing expression of luciferase and/or brat-dsRNA and/or Spt5-siRNA in NB II. ***B***, adult brains were stained for the synaptic protein Bruchpilot (green) to label neuropil structures and the nuclear membrane protein Lamin (red) to visualize the brain cortex. Single pictures were taken at the level of the ellipsoid body of the central complex. Scale bar: 50 µm. ***C***, quantification of luciferase activity, relative to that of control flies; n=6-16 single adult flies per genotype. ***D***, NB II lineages in brains from 3rd instar larvae were marked with mCD8::GFP (green) and co-stained for the nuclear proteins Dpn and Ase to distinguish the large NB II (Dpn+ Ase-), newborn INPs (Dpn- Ase-), immature INPs (Dpn- Ase+), and mature INPs (Dpn+ Ase+). Neighboring type I NBs co-express Dpn and Ase. In control brains, two out of eight NB II lineages per brain hemisphere are shown. Spt5-knockdown causes incomplete NB II lineages, whereas brat-knockdown results in massive expansion of cells with characteristics of NB II (Dpn+ Ase-). In the double-knockdown, separate clusters like in controls are observed, but cells maintain mostly NB II characteristics and only few cells express Ase as an indicator of further differentiation. Scale bar: 10 µm. ***E***, EdU incorporation (cyan) in S-phase cells within a period of 90 min in whole mount brain preparations. Compact EdU signals are seen in the lateral regions representing the proliferation centers of the optic lobes, dispersed signals are evident in the central brain with NB II and their lineages labeled in green. Scale bar: 100 µm. ***F***, in higher magnifications, many proliferating cells outside and within NB II lineages (outlined with dashed lines) are seen in controls, with a strong increase upon brat-KD. No EdU positive cells are detected in NB II lineages under Spt5-KD and brat-KD/Spt5-KD conditions. Scale bar: 10 µm.

To study the underlying cellular differences between the different genotypes, we analyzed NB II lineages in 3^rd^ instar larval brains by concurrent expression of GFP and stainings for Deadpan (Dpn) and Asense (Ase). In control flies, there are only 8 NB II in each brain hemisphere, which express Dpn but not Ase (Dpn+ Ase-), in contrast to approximately 100 type I neuroblasts, where both proteins are present (Dpn+ Ase+). The intermediate neural progenitors (INPs) generated by each NB II pass through a maturation process (from Dpn- Ase-, to Dpn-Ase+, to Dpn+ Ase+) before ganglion mother cells are born (Fig. 3D). As reported previously, brat-knockdown causes a massive expansion of NB II like cells (Dpn+ Ase-) at the expense of INPs (Bowman *et al*., 2008; Janssens *et al*., 2014; Komori *et al*., 2018; Xiao *et al*., 2012). Brain hemispheres were enlarged, with the dorsal part being nearly completely covered with Dpn+ Ase-cells without signs of further lineage progression (Fig. 3D). Spt5-knockdown resulted in a strong suppression of the overgrowth phenotype caused by brat-knockdown and reduced the total number of cells within each lineage, but nevertheless allowed the generation of mature INPs (Ase+ Dpn+) (Fig. 3D). Distinct GFP-positive cell clusters were visible similar to the control situation. However, within each cluster most cells still displayed NB II characteristics (Dpn+ Ase-) and only few cells expressed Ase as a marker for INP maturation (Fig. 3D). Based on these observations we concluded that, although Spt5-knockdown cannot efficiently revert transformed NB II like cells into further differentiated INPs, it nevertheless has a major negative impact on tumor formation, possibly by interfering with NB II proliferation. We confirmed this assumption by pulse labeling S-phase cells with EdU in larval brains (Figs. 3E-F). Knockdown of Spt5 alone or in combination with brat abolished EdU-incorporation within the GFP-labeled cell clones (highlighted areas), whereas the brains with brat-knockdown alone contained multiple cells in S-phase that actively incorporated EdU. Although we noticed a moderate increase in apoptotic cells (positive for the cleaved effector caspase Dcp-1) in brat-*/*Spt5-knockdown conditions within the GFP-labeled cell clones (Fig. S3B), the major tumor suppressive mechanism of Spt5-knockdown can be ascribed to impaired proliferation.

### Effects of Spt5 on tumor transcriptomes

To identify the molecular basis of the observations described above, we isolated NB II from 96 hours-old larvae and analyzed their transcriptomes. As shown in Fig. 4A, control and brat-/Spt5-codepleted cells were highly similar to each other and clearly distinct from brat-depleted (tumorous) cells with respect to principal component 1, consistent with the reversion of overgrowth by Spt5-knockdown.

**Figure 4.**
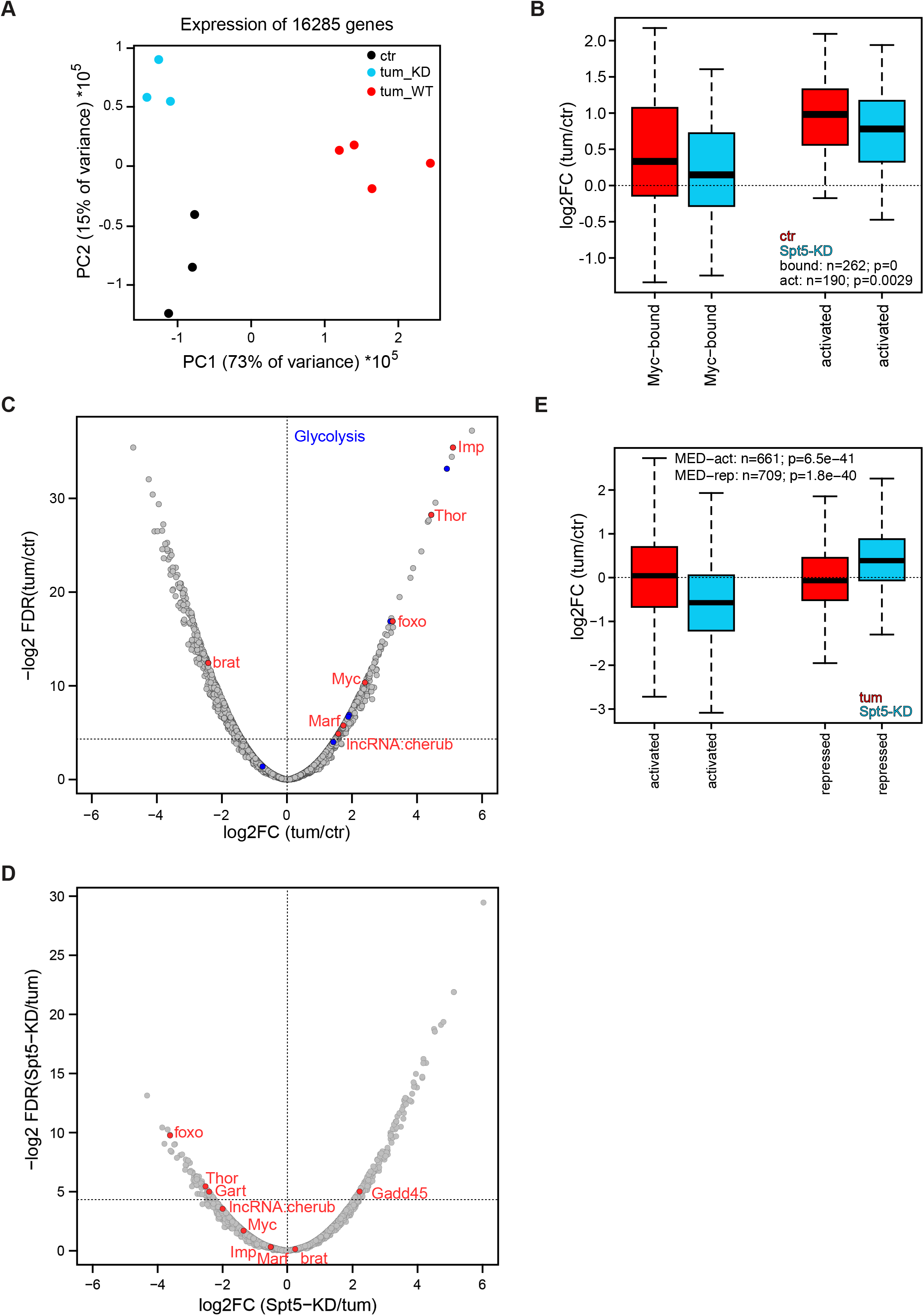
Effects of brat- and Spt5-knockdown on NB II transcriptomes. ***A***, principal component analysis of NB II transcriptomes from control (black, “ctr”), tumorous (red, “tum_WT”) or tumorous flies with Spt5-knockdown (blue, “tum_KD”). ***B***, expression levels of Myc-bound or -activated genes that were previously identified in cultured S2 cells (Herter *et al*., 2015) in brat-depleted NB II relative to control NB II (red), and in Spt5-/brat-codepleted NB II relative to control NB II (blue). ***C, D***, volcano plots showing expression in brat-depleted NB II (tumors) relative to control NB II (***C***), and in Spt5-/brat-codepleted NB II relative to brat-depleted NB II (***D***). Horizontal lines mark significance level (FDR q-value) of 0.05. Labeled genes are described in the text; for a complete listing of all genes see Tables S2,3. ***E***, expression levels of previously identified Med27-activated or -repressed genes in brat-depleted NB II relative to control NB II (red), and in Spt5-/brat-codepleted NB II relative to control NB II (blue). P values are derived from a paired Student’s t-test.

Comparison of control with brat-depleted neuroblasts revealed several alterations of uncharacterized genes (shown in grey) as well as expected changes in gene expression (Figs. 4B,C): brat levels were clearly reduced, whereas the RNA-binding protein Imp (Samuels *et al*, 2020), the long noncoding RNA cherub (Landskron *et al*, 2018), the mitochondrial fusion factor Marf (Bonnay *et al*, 2020), Myc and Myc target genes (Betschinger *et al*., 2006; Herter *et al*, 2015; Neumuller *et al*, 2013), as well as glycolytic enzymes (Bonnay *et al*., 2020; van den Ameele & Brand, 2019) were all strongly upregulated in response to brat-knockdown. All of these changes had been observed before and they contribute to the tumorous phenotype. In addition, the transcription factor Foxo and its target Thor/4E-BP were overexpressed in brat-depleted NB II.

Next, we analyzed the impact of Spt5-knockdown on tumors caused by brat-knockdown. Brat levels themselves were not altered, but Myc targets were significantly down-regulated, in line with observations in mammalian cancer cells (Fig 4B,D). The other described genes were moderately (Marf, Imp) or strongly (lncRNA:cherub) reduced in their expression upon Spt5-knockdown (Figs. 4D). In addition, Gart (the second enzyme of the purine biosynthesis pathway; Welin *et al*, 2010) was significantly repressed, and Gadd45 (an inhibitor of cell cycle progression and inducer of apoptosis; Tamura *et al*, 2012) was strongly activated. We also noted that the levels of Foxo and Thor/4E-BP dropped in Spt5-knockdown cells. Together, these expression changes are sufficient to explain the reduction in tumor growth and cellular proliferation and most of them can be ascribed to an impairment of Myc-dependent gene activation upon Spt5-knockdown. However, some of the affected genes are not *bona fide* Myc targets, e.g. lncRNA:cherub (Herter *et al*., 2015). To find other candidate upstream regulators of these genes, we explored publicly available NB II transcriptome data, and found a significant correlation between Spt5-controlled genes and Mediator target genes. Notably, Gart, lncRNA:cherub, Foxo, and Thor all require Mediator for their full expression (Fig 4E; Homem *et al*, 2014), raising the possibility that Spt5 might affect their expression via an interaction with Mediator.

### Organismal consequences of Spt5 depletion

Despite the massive brain overgrowth upon brat-knockdown in NB II lineages, the tumor-bearing animals reached adulthood at expected frequencies (Fig. S4A). However, all of them died within less than 10 days of eclosion, whereas the majority of control flies were still alive after 60 days (Fig. 5A; for statistical significance of various comparisons see Table S1). Myc-knockdown slightly extended the survival of tumor-bearing flies, showing that these tumors are Myc-dependent (Fig. S4B). The survival benefits are presumably limited by the requirement for Myc in NB II, as seen by the reduced longevity upon single Myc-knockdown (Fig. S4B). In contrast, Spt5-knockdown did not impair the survival of control flies, but extended the life span of tumor-bearing animals to more than 26 days (Fig. 5A). This rescue was fully reverted by co-expression of an siRNA-resistant version of Spt5, ruling out off-target effects. Overexpression of Spt5 on its own had the opposite effect and significantly shortened the life of tumorous animals, but had only minor effects on healthy controls. Together, these observations emphasize the importance of Spt5 for abnormal, tumorous tissue growth.

**Figure 5.**
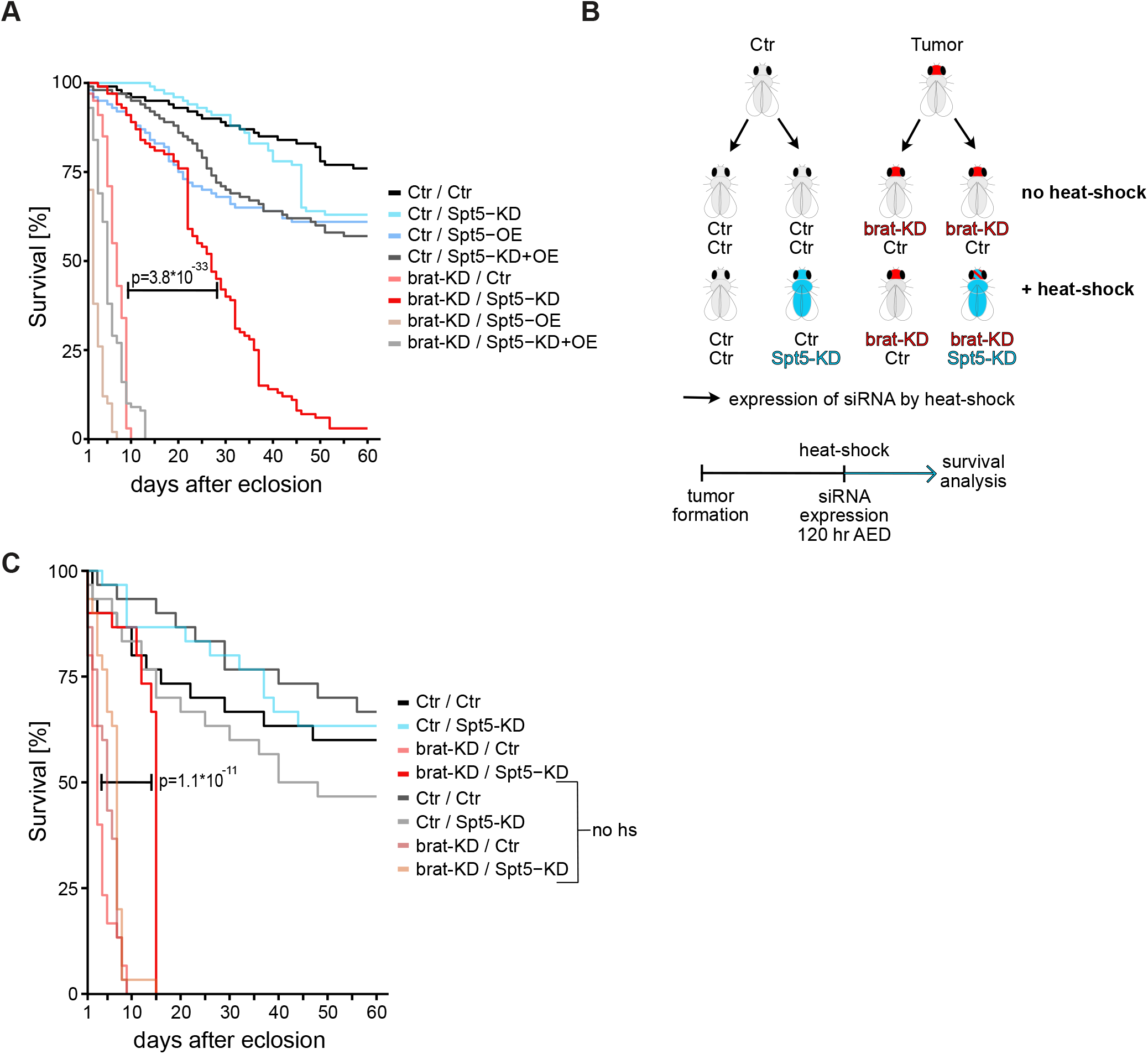
Impact of Spt5-knockdown on longevity of tumorous flies. ***A***, survival of male flies with the indicated genotypes in days after adult eclosion (n=100 flies for each genotype). ***B***, scheme for ubiquitous and temporally controlled Spt5-depletion in tumorous and control animals (for details see text). ***C***, survival of male flies with the indicated genotypes +/-heat-shock induced ubiquitous Spt5-depletion in days after adult eclosion. Spt5-knockdown significantly extended the lifespan of tumorous flies (p=1.1 * 10^-11^; n=30 flies for each genotype).

To explore whether this dependency could potentially be exploited in a curative context, we modified the NB II tumor model. In this new setup, NB II tumors are induced with the same brat-knockdown transgene as used above. In contrast, Spt5-knockdown is driven by the Actin5C promoter that is ubiquitously active in the entire organism. This transgene is initially activated by a heat-shock, administered to 120 hours-old larvae (well after the onset of GAL4-expression driving brat-knockdown in NB II; Albertson *et al*, 2004) and remains active thereafter (Fig. 5B). The transgene expresses the same Spt5-siRNA as used in the earlier setup, although presumably at a lower level, since this approach does not involve any GAL4/UAS amplification loop. When flies carrying brat- and Spt5-knockdown transgenes were reared in the absence of heat-shock, they succumb to tumors within 10 days of adult eclosion; control flies lacking the brat-knockdown transgene had the expected life span (Fig. 5C). After heat-shock, flies lacking the siSpt5-transgene showed an analogous behavior. However, in combination with heat-shock the siSpt5-transgene almost doubled the lifespan of tumorous flies (Fig. 5C; Table S1). We conclude that systemic targeting of Spt5 is beneficial for cancer bearing flies.

## Discussion

Several experimental approaches allow the identification of potential cancer drug targets at a medium-to large-scale level. These include the analysis of gain-of-function or overexpression mutations in human tumor samples, systematic knockdown or knockout screens in human cancer cell lines (e.g. Boehm & Golub, 2015), silencing or depletion of candidate genes in mouse transplant models. However, targets identified by these approaches could also be relevant for healthy tissues. It is therefore essential to determine the “therapeutic window” of any putative target. This is usually done by analysing appropriate mouse models, containing e.g. floxed target genes in combination with an OHT-activatable Cre recombinase, or expressing shRNAs against the target gene. Such approaches are more laborious and expensive than the initial genetic screens, and hence therapeutic windows are often addressed only once target-specific inhibitors are available, resulting in high attrition rates at late pre-clinical stages. Our present analysis suggests that Drosophila can be used to reveal the existence of such therapeutic windows.

The elongation factor Spt5 initially caught our attention because of its physical interaction with Myc in cultured human cancer cells (Baluapuri *et al*., 2019). Here, we found that it also collaborates with Myc functionally in vivo. Simultaneous reduction of both proteins during larval development synergistically impaired the growth of imaginal tissue, consistent with the notion that Myc-dependent efficient activation of growth-promoting genes requires association with Spt5. Combining Spt5-knockdown with Myc-overexpression during post-proliferative eye disc development resulted in a striking novel phenotype, indicative of massive apoptosis not seen with either treatment alone. This could indicate that some Myc targets do not require Spt5 for their expression, and that the balance of Spt5-dependent and -independent targets determines the biological outcome of Myc activation, e.g. tissue growth versus attrition (similar to what was suggested by Steiger *et al*, 2008). Alternatively, combined Spt5-knockdown and Myc-overexpression might titrate Spt5 away from some genes, affecting their expression and resulting in the observed phenotype (similar to what was suggested by Baluapuri *et al*., 2019).

We used Spt5 as an example of an essential Myc co-factor and evaluated the consequences of knocking down Spt5 in a Myc-dependent NB II brain tumor model. In a first approach, we used the same expression system to target both brat (in order to generate the NB II tumors) and Spt5 specifically in NB II. In this setting, Spt5-knockdown almost completely reverted the tumorous tissue overgrowth and more than tripled adult animal survival. Knockdown of Spt5 in selected neuroblasts of control animals without brain tumors had mild effects on brain development, and did not negatively impact adult survival, demonstrating the potential value of Spt5 as a therapeutic target. However, in clinical settings it is typically not possible to direct a therapy exclusively at transformed cells and therapeutic intervention cannot be initiated at early tumor development. For this reason, we developed a second system that allowed temporal separation of tumor initiation and Spt5-knockdown, and that targeted Spt5 not only in NB II but throughout the organism. While this approach relied on the same system to deplete brat and the same Spt5-targeting siRNA as the first approach above, the latter was induced by a heat-shock and directly driven by the Actin5C promoter rather than being amplified by a GAL4/UAS loop, presumably resulting in lower siRNA expression and less efficient Spt5 depletion in NB II. Nevertheless, Spt5-knockdown had a strong therapeutic benefit for tumorous flies, as it almost doubled their survival time. Importantly, ubiquitous Spt5-knockdown did not impair the survival of tumor-free control animals, nor did heat-shocks per se have any deleterious effect on longevity. A molecular explanation for this tumor-suppressive effect is provided by our analysis of NB II transcriptomes: Spt5-knockdown resulted in strong down-regulation of several genes associated with NB II transformation, and an up-regulation of genes opposing uncontrolled proliferation. Most of these expression changes can be ascribed to a reduction of Myc:Spt5 complexes, while some probably reflect a functional interaction of Spt5 with the Mediator complex, which itself plays a role in NB II tumor formation. As expected, Myc-knockdown also extended the longevity of tumor-bearing flies, but this effect was less pronounced than for Spt5-knockdown. This difference might indicate that Myc:Spt5 complexes are more critical for transformed cells than for normal tissues. In any case, our experiments demonstrate that targeting a protein, Spt5, which was selected based on its physical interaction with Myc, can reduce tumor mass and provide a survival benefit for tumor-bearing animals, even though this protein is essential for normal development. It is open, though, whether Spt5 is the best-suited target in Myc-dependent cancers, as many additional proteins have been shown to bind Myc. Our Drosophila-based approach allows a simple pre-screening of these candidates to filter for the best targets that can subsequently be funneled into more laborious analyses in mice.

### Author contributions

JH and AO performed the cloning and RNAseq experiment. PG analysed the RNAseq data. AO, AK and LMB conducted the in vivo experiments. AO, LMB and TR performed microscopy. TR and PG planned the in vivo experiments. JH, EW, TR and PG designed the study, supervised the experiments and wrote the paper.

## Acknowledgements

We thank David Gilmour, Erich Buchner, Fernando Diaz-Benjumea, Jürgen Knoblich and Georg Krohne for generously providing antibodies and fly stocks, Hugo Stocker for critical reading of the manuscript, the DFG for grants WO 2108/1-1 & GRK 2243 to EW, and the ERC for grant TarMyc to EW.

---

**Figure S1:**
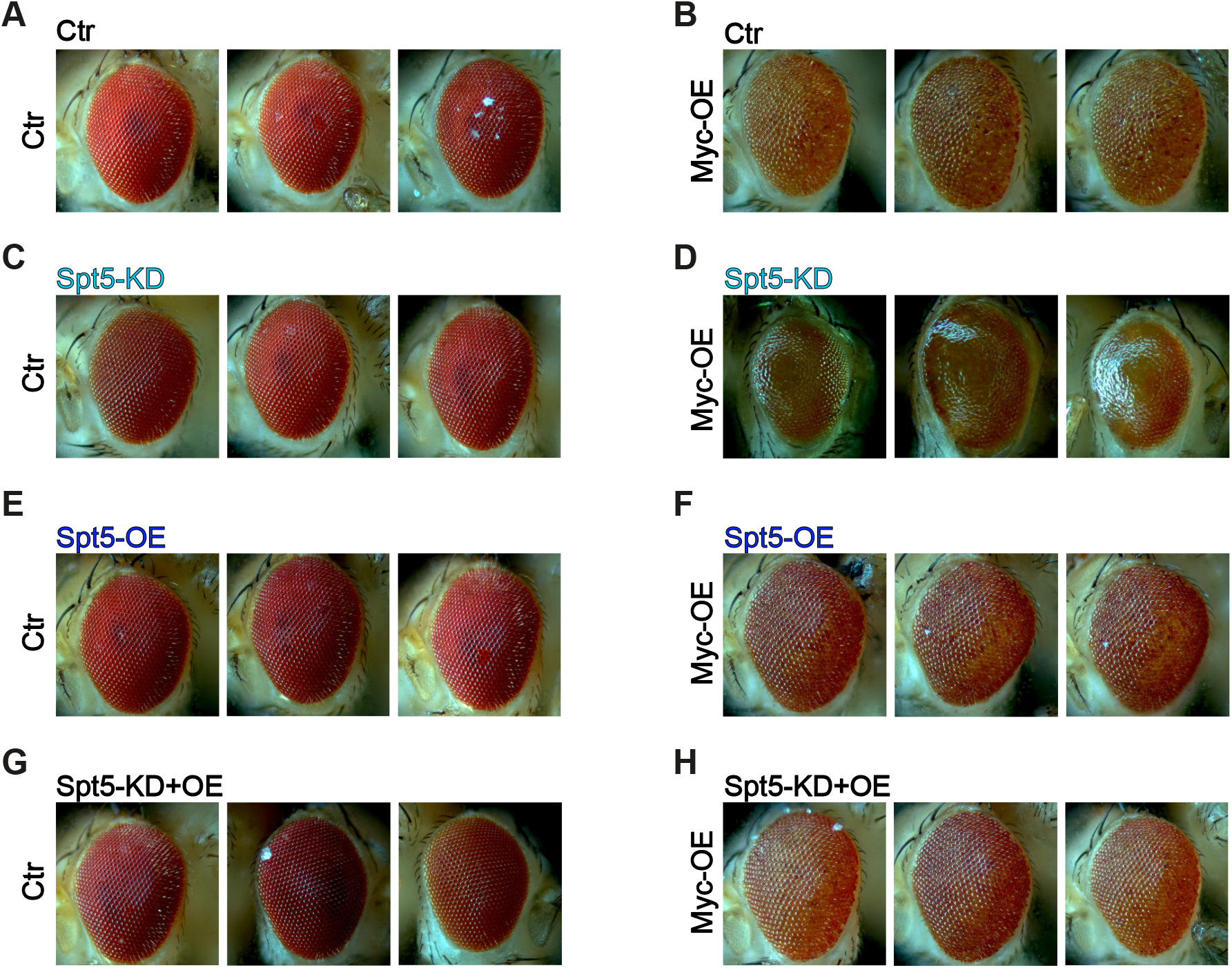
***A-H***, pictures of adult eyes having experienced GMR-driven Myc-overexpression +/-Spt5-overexpression +/- Spt5-knockdown. Pertains to Fig. 1C,D.

**Figure S2:**
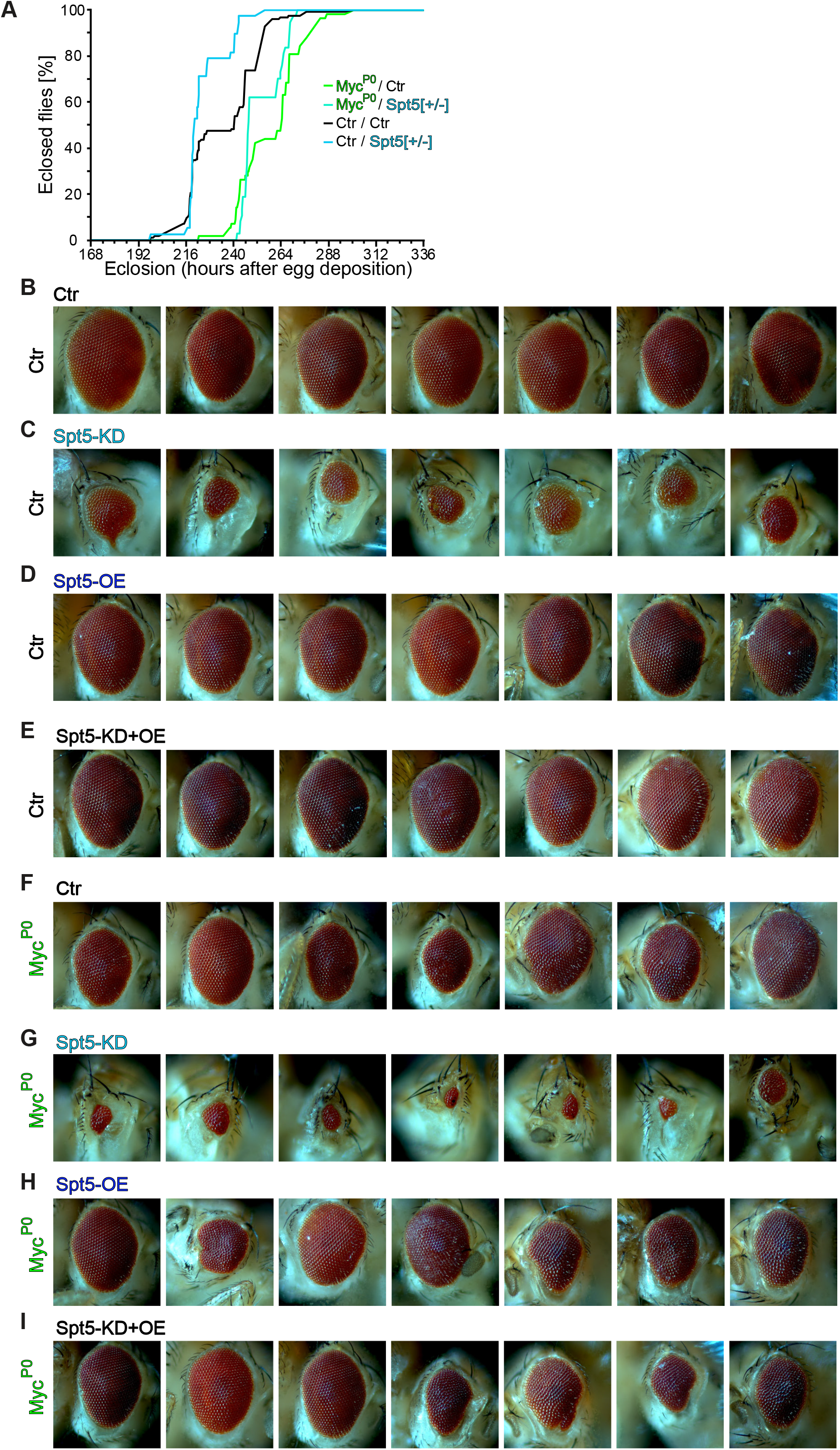
***A,*** duration of development, in hours from egg deposition to adult eclosion (n=34-82 males per genotype). ***B-I,*** pictures of adult eyes having experienced Spt5-overexpression and/or Spt5-knockdown in the background of wildtype or mutant Myc. Pertains to Fig. 2 C,D.

**Figure S3:**
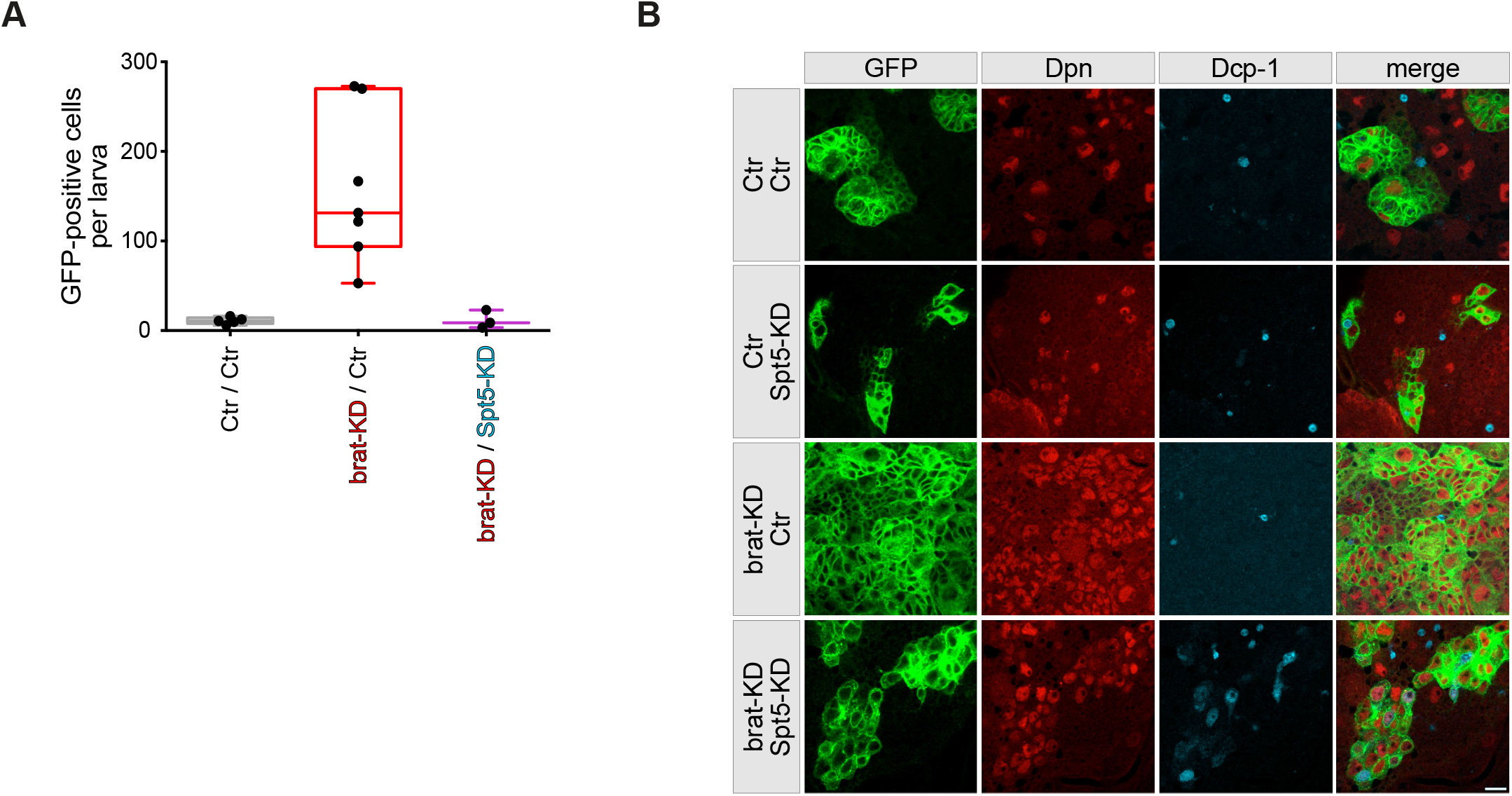
***A***, number of NB II-derived (GFP-positive) cells per larva (based on 3-7 independent dissections per genotype, each involving 130-240 larvae). ***B***, NB II lineages in brains from 3rd instar larvae were marked with mCD8::GFP (green) and co-stained for the nuclear proteins Dpn (red) and the apoptosis marker Dcp-1 (blue). Few apoptotic cells are observed in control, Spt5-KD and brat-KD brains but their number increases under double-knockdown conditions. Scale bar: 10 µm.

**Figure S4:**
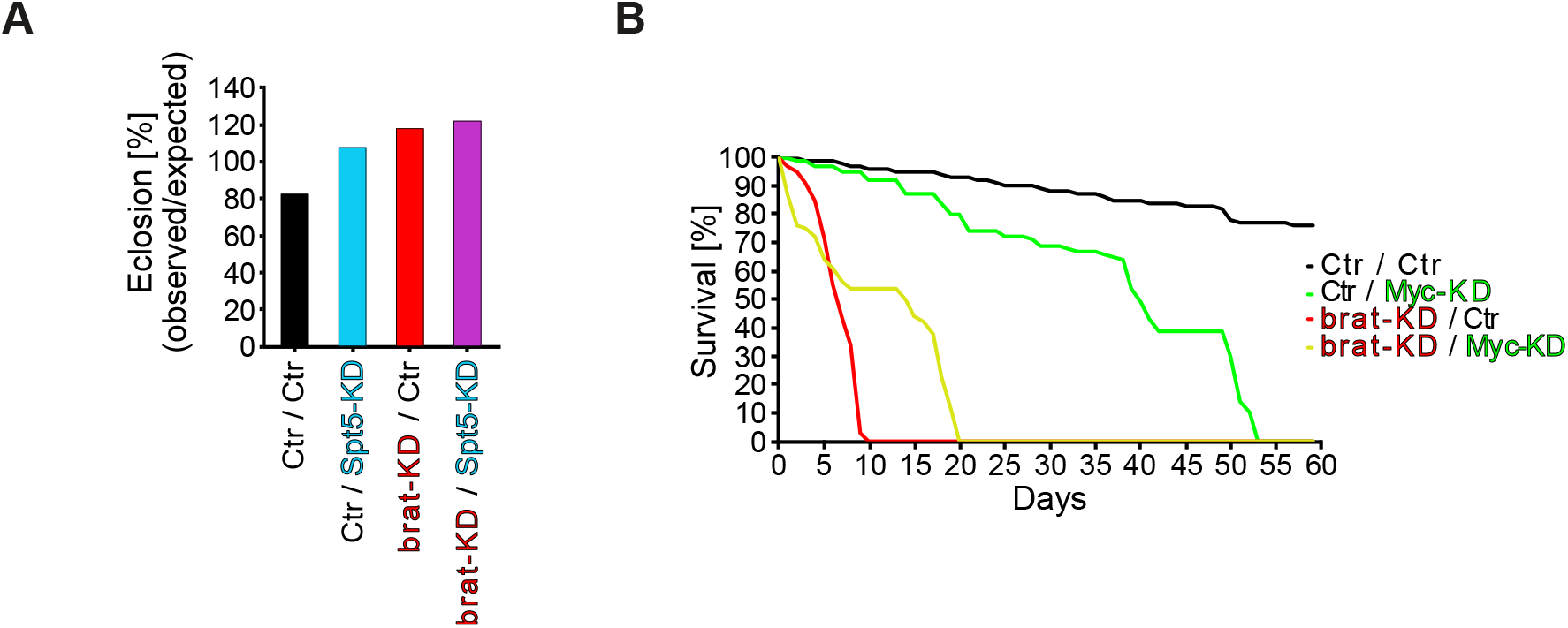
***A***, eclosion rate of flies with or without NB II tumors in the presence or absence of Spt5-knockdown. Percentage of eclosed adult flies relative to the expected fraction (n=171-271 per genotype). ***B***, effect of Myc-depletion in tumorous or normal NB II on survival of adult male flies (n=65 to 100 flies for each genotype).

**Table S1:** statistical significance of differences in longevity. Pertains to Fig. 5A,C.

**Table S2:** raw read counts from control NB II or brat-knockdown NB II +/- Spt5-knockdown. Pertains to Fig. 4B,C.

**Table S3:** normalized expression values, p-values & FDR q-values from pairwise comparisons of genotypes. Pertains to Fig. 4B,C.

## Materials & Methods

### Flies

Sources of flies: “GMR-GAL4” and “GMR-GAL4 3x(UAS-Myc)” were characterized by (Montero *et al*, 2008; Steiger *et al*., 2008); “wor-GAL4 ase-GAL80 UAS-GFP UAS-Luciferase” and “wor-GAL4 ase-GAL80 UAS-brat-IR UAS-GFP UAS-Luciferase” were initially generated by (Neumuller et al., 2013) and also described in (Herter et al., 2015); “ey-FLP tub-FRT-Myc-FRT-GAL4” was described in (Bellosta et al., 2005); UAS-Spt5[resistant to siSpt5] (Qiu & Gilmour, 2017); UAS-siSpt5 (Bloomington stock number B-34837; Perkins *et al*, 2015); mutant allele “Spt5[SIE-27]“ (Mahoney *et al*., 2006). “act-FRT-stop-FRT-siSpt5” was generated by inserting “AggccagtCAGAAGCTACAGTCCATTCAAtagttatattcaagcataTTGAATGGACTGTAGCTTCTGgcggccAGTC” (“siSpt5_f”) into pAct-FRT-stop-FRT3-FRT-FRT3-GAL4_attB (AddGene vector #52889; Bosch *et al*, 2015). The resulting construct pACT5C-FRT-stop-FRT-siSpt5 was inserted in ZH86Fb by GenetiVision Corp (Houston, Tx).

### Genetic manipulation of Spt5 and Myc in eyes

Eye specific reduction of Myc levels as used for Fig. 2C,D was described by Bellosta et al. 2005. Briefly, a Myc cDNA was ubiquitously expressed under the control of the tubulin promoter by the transgene “tub-FRT-Myc-FRT-GAL4” (inserted on the X-chromosome), which increases Myc levels to <180% as compared to control (Wu & Johnston, 2010). The same X-chromosome carries an ey-FLP transgene, which eliminates the Myc cDNA specifically in eye imaginal disc cells, resulting in expression of GAL4 instead. Flies designated as “Myc^P0^” additionally carry the hypomorphic allele *Myc^P0^* on the same X-chromosome, whereas “ctr” flies are wild type for Myc and only carry the two described transgenes. Hence, eye imaginal discs of the Myc^P0^ flies described in Fig. 1C,D are mutant for Myc specifically in the eye primordia, thus expressing less than 40% of Myc mRNA. Importantly, this Myc allele only reduces the amount of Myc protein, but does not alter its amino acid sequence.

### Targeted expression

Type II neuroblasts were targeted by a combination of worniu (wor)-GAL4, which is expressed in type I and II NBs, and asense (ase)-GAL80 to repress GAL4 activity in the type I NBs (Neumüller *et al*, 2011).

To knock down Spt5 ubiquitously after the onset of tumor generation, the system above (wor-GAL4 ase-GAL80 UAS-brat-KD) was combined with the transgenes “hs-FLP” and “pACT5C-FRT-stop-FRT-siSpt5”. siSpt5 expression was initiated by transferring larvae at 120 h after egg deposition for 1 h to a water bath at 37°C.

### Relevant genotypes

**Figure 1C-D; Figure S1**

GMR-GAL4/+

GMR-GAL4/+; UAS-siSpt5/+

GMR-GAL4/+; UAS-Spt5/+

GMR-GAL4/+; UAS-Spt5 UAS-siSpt5/+

GMR-GAL4 3x(UAS-Myc)/+

GMR-GAL4 3x(UAS-Myc)/+ ; UAS-siSpt5/+

GMR-GAL4 3x(UAS-Myc)/+ ; UAS-Spt5/+

GMR-GAL4 3x(UAS-Myc)/+ ; UAS-Spt5 UAS-siSpt5/+

**Figure 2A ; Figure S2A**

+/Y

+/Y; Spt5[SIE-27]/+

dm[P0]/Y

dm[P0]/Y; Spt5[SIE-27]/+

**Figure 2C-D**; **Figure S2B-I**

tub-FRT-Myc-FRT-GAL4 ey-FLP/Y

tub-FRT-Myc-FRT-GAL4 ey-FLP/Y; UAS-siSpt5/+

tub-FRT-Myc-FRT-GAL4 ey-FLP/Y; UAS-Spt5/+

tub-FRT-Myc-FRT-GAL4 ey-FLP/Y; UAS-Spt5 UAS-siSpt5/+

dm[P0] tub-FRT-Myc-FRT-GAL4 ey-FLP/Y

dm[P0] tub-FRT-Myc-FRT-GAL4 ey-FLP/Y; UAS-siSpt5/+ dm[P0]

tub-FRT-Myc-FRT-GAL4 ey-FLP/Y; UAS-Spt5/+

dm[P0] tub-FRT-Myc-FRT-GAL4 ey-FLP/Y; UAS-Spt5 UAS-siSpt5/+

**Figure 3B,D,E**; **Figure 4**; **Figure S3**; **Figure S4A**

wor-GAL4 ase-GAL80 UAS-mCD8::GFP

wor-GAL4 ase-GAL80 UAS-mCD8::GFP UAS-brat-KD

wor-GAL4 ase-GAL80 UAS-mCD8::GFP UAS-siSpt5

wor-GAL4 ase-GAL80 UAS-mCD8::GFP UAS-brat-KD UAS-siSpt5

**Figure 3C**

wor-GAL4 ase-GAL80 UAS-RLuc

wor-GAL4 ase-GAL80 UAS-RLuc UAS-siSpt5

wor-GAL4 ase-GAL80 UAS-RLuc UAS-Spt5

wor-GAL4 ase-GAL80 UAS-RLuc UAS-siSpt5 UAS-Spt5

wor-GAL4 ase-GAL80 UAS-FLuc UAS-brat-KD

wor-GAL4 ase-GAL80 UAS-FLuc UAS-brat-KD UAS-siSpt5

wor-GAL4 ase-GAL80 UAS-FLuc UAS-brat-KD UAS-Spt5

wor-GAL4 ase-GAL80 UAS-FLuc UAS-brat-KD UAS-siSpt5 UAS-Spt5

**Figure 5A**

wor-GAL4 ase-GAL80 UAS-RLuc

wor-GAL4 ase-GAL80 UAS-RLuc UAS-siSpt5 wor-GAL4 ase-GAL80 UAS-RLuc UAS-Spt5

wor-GAL4 ase-GAL80 UAS-RLuc UAS-siSpt5 UAS-Spt5

**Figure 5C**

hs-FLP wor-GAL4 ase-GAL80

hs-FLP wor-GAL4 ase-GAL80 UAS-brat-KD

hs-FLP wor-GAL4 ase-GAL80 act-FRT-stop-FRT-siSpt5

hs-FLP wor-GAL4 ase-GAL80 UAS-brat-KD act-FRT-stop-FRT-siSpt5

**Figure S4B**

wor-GAL4 ase-GAL80

wor-GAL4 ase-GAL80 UAS-brat-KD

wor-GAL4 ase-GAL80 UAS-Myc-KD

wor-GAL4 ase-GAL80 UAS-brat-KD UAS-Myc-KD

### Confocal microscopy

For immunostainings, brains from late 3rd instar larvae or adults were dissected in PBS (10 mM Na_2_HPO_4_, 2 mM KH_2_PO_4_, 2.7 mM KCl, 137 mM NaCl) and fixed on ice for 25 min in PLP solution (4% paraformaldehyde, 10 mM NaIO_4_, 75mM lysine, 30 mM sodium phosphate buffer, pH 6.8). All washings were done in PBT (PBS plus 0.3% Triton X-100). After blocking in PBT containing 5% normal goat serum for 1 h, tissues were incubated overnight at 4°C with combinations of the following primary antibodies: rabbit anti-Ase (1:400; F. Diaz-Benjumea, Madrid, Spain), mouse anti-Bruchpilot (1:30, clone nc82; E. Buchner, Würzburg, Germany), rabbit anti-Dcp-1 (1:100, Cell Signaling Techn. # 9578, Danvers, MA, USA), guinea pig anti-Dpn (1:1000, J. Knoblich, Vienna, Austria), chicken anti-GFP (1:1500; abcam #ab13970, Cambridge, UK), guinea pig anti-Lamin DmO (1:300; G. Krohne, Würzburg, Germany). Secondary antibodies conjugated with AlexaFluor 488, Cy3 or Cy5-conjugated were purchased from Molecular Probes (Eugene, OR, USA) and Dianova (Hamburg, Germany).

For 5-ethynyl-2’-deoxyuridine (EdU) labeling, larval brains from 3rd instar larvae were dissected in PBS and incubated with 20 µM EdU in PBS for 90 min. After fixation in 4% paraformaldehyde for 15 min, followed by immunostaining for GFP and Dpn, before EdU incorporation into replicating DNA was detected with the Click-iT® Alexa Fluor 647 EdU imaging kit (Thermo Fisher Scientific (Invitrogen), Waltham MA, USA). Embedding of brains was done in Vectashield (Vector Laboratories, Burlingame, CA, USA) and confocal images were collected with a Leica SPE or SP8 microscope (Leica Microsystems, Wetzlar, Germany). Image processing was carried out with the ImageJ distribution Fiji (Schindelin *et al*, 2012).

### Phenotypic Analysis

To measure adult eye sizes, adult males were collected at 1 to 7 days after eclosion and killed by freezing. One eye per individual fly was photographed on a Zeiss Discovery.V8 stereomicroscope fitted with a 1.5x lens and processed with Axiovision Extended Focus software and the ImageJ distribution Fiji.

To measure luciferase activity, male flies were collected within one day of adult eclosion and frozen individually at -20°C until use. Each fly was then lysed in 50 µl Passive Lysis Buffer (Promega) and homogenized with approximately 10 steel beads in a ‘Bullet Blender Blue’ Homogenizer at speed 10 for 2 minutes, followed by a 4’ centrifugation at 12,000 g. Ten μl of the supernatant was transferred into a black 96-well plate and assayed for luciferase expression using the Dual Luciferase Reporter Assay System in an automated luminometer. Note that the tumorous brat-KD flies contain a “UAS-FireflyLuciferase” transgene, whereas the non-tumorous flies without the brat-KD carry a “UAS-RenillaLuciferase” transgene. Hence, luciferase activities can only be compared within each series of genotypes, but not between the brat-WT genotypes (shown in black in Fig 3C) and the brat-KD genotypes (shown in red in Fig 3C).

For weighing flies, 1 to 4 day old adult flies were dried for 20’ at 95° (first for 10’ with a closed, then with an opened lid) and then stored at room temperature. Before weighing on a Mettler UMT5 Comparator scale (Mettler Toledo), the flies were allowed to equilibrate with ambient atmosphere for at least 30’.

To determine duration of development, timed egg lays (5 – 14 h) were performed and eclosion was monitored 2 to 3 times a day.

### Survival analysis

Parents were transferred to a fresh food vial every three days. Offspring was collected within one day of adult eclosion, and subsequently transferred to fresh vials every two days. The number of living flies was monitored daily for a period of 60 days.

### Isolation of type II neuroblasts

Processing of larvae for next-generation sequencing was carried out as described by (Harzer *et al*, 2013). Briefly, five-day old larvae were washed sequentially in PBS, 70% ethanol and Schneider’s medium. Within ≤1 h larvae were dissected and brains transferred to a 0.5 ml low-binding Eppendorf tube containing Rinaldini’s solution (8 g/l NaCl, 0.2 g/l KCl, 50 mg NaH_2_PO_4_, 1 g/l NaHCO_3_, 1 g/l Glucose). After 2 washes, Rinaldini’s solution was replaced with dissociation solution (Schneider’s medium containing 100 ml/l heat-inactivated fetal bovine serum, 2 ml/l insulin, 20 ml/l penicillin-streptomycin, 100 ml L-glutamine, 20 mg/l L-glutathione, 20 mg/ml collagenase I, 20 mg/ml papain), and the brains were stirred up by pipetting. After one hour incubation at 30°C with occasional mixing, the brains were washed twice with Rinaldini’s solution and with Schneider’s medium, and then mechanically dissociated by pipetting. The resulting cell suspension was filtered through a 30-µm mesh 5-ml FACS tube, which was then filled up with Schneider’s medium to a total volume of 10 µl per dissected larval brain and sorted in a BD FACSAria™ III sorter. Type II Neuroblasts were identifed based on side scatter (SSC), forward scatter (FSC) and GFP intensity, collected into 96-well microtiter plates, containing 1 μl β-mercaptoethanol and 100 μl Lysis Buffer (Agilent Technologies Absolutely RNA Nanoprep Kit) per well, and subsequently stored at −80 °C until use.

### mRNA library preparation

RNA was isolated using Agilent Technologies’ Absolutely RNA Nanoprep Kit (including DNase I digestion). RNA concentration and quality were determined on 2100 Bioanalyzer Instrument (Agilent Technologies) using the Agilent RNA 6000 Pico Kit (Agilent Technologies). Library preparation was performed using the Poly(A) mRNA Magnetic Isolation Module (New England Biolabs) and the NEBNext Ultra II Directional RNA library Prep Kit for Illumina (New England Biolabs). For library amplification 17 or 24 PCR cycles were used. Library size distribution and concentration were analyzed on the Fragment Analyzer (Agilent Technologies) using the NGS Fragment High Sensitivity Analysis Kit (1-6,000 bp; Agilent Technologies). The libraries were sequenced on Illumina instrument (NEXTSeq500).

### Bioinformatics

Bliss synergy scores (Bliss, 1939) were calculated using the R package synergyfinder 1.10.7 (Zheng *et al*, 2022), where scores >10 suggest a synergistic interaction; n=6 to 10 collections per genotype for Fig. 2A, median derived of 8 flies for each genotype for Fig. 2C,D.

For RNAseq analysis, reads were mapped to version BDGP6 of the *Drosophila* genome, using bowtie2 with the setting “very-sensitive-local” (Langmead & Salzberg, 2012) (2.2 to 9.8 million mapped reads per sample). Differentially expressed genes were identified using edgeR 3.26.8 (Robinson *et al*, 2010). Gene set enrichment analysis was carried out with GSEA 4.0.2. (Subramanian *et al*, 2005) and GO-terms obtained from the ENSEMBL annotation for BDGP6.32. Volcano & box plots were generated in R.

